# Myosin-binding protein C forms C-links and stabilizes OFF states of myosin

**DOI:** 10.1101/2023.09.10.556972

**Authors:** Anthony L. Hessel, Nichlas M. Engels, Michel Kuehn, Devin Nissen, Rachel L. Sadler, Weikang Ma, Thomas C. Irving, Wolfgang A. Linke, Samantha P. Harris

## Abstract

Contraction force in muscle is produced by the interaction of myosin motors in the thick filaments and actin in the thin filaments and is fine-tuned by other proteins such as myosin-binding protein C (MyBP-C). One form of control is through the regulation of myosin heads between an ON and OFF state in passive sarcomeres, which leads to their ability or inability to interact with the thin filaments during contraction, respectively. MyBP-C is a flexible and long protein that is tightly bound to the thick filament at its C-terminal end but may be loosely bound at its middle- and N-terminal end (MyBP-C^C1C7^). Under considerable debate is whether the MyBP-C^C1C7^ domains directly regulate myosin head ON/OFF states, and/or link thin filaments (“C-links”). Here, we used a combination of mechanics and small-angle X-ray diffraction to study the immediate and selective removal of the MyBP-C^C1C7^ domains of fast MyBP-C in permeabilized skeletal muscle. After cleavage, the thin filaments were significantly shorter, a result consistent with direct interactions of MyBP-C with thin filaments thus confirming C-links. Ca^2+^ sensitivity was reduced at shorter sarcomere lengths, and crossbridge kinetics were increased across sarcomere lengths at submaximal activation levels, demonstrating a role in crossbridge kinetics. Structural signatures of the thick filaments suggest that cleavage also shifted myosin heads towards the ON state – a marker that typically indicates increased Ca^2+^ sensitivity but that may account for increased crossbridge kinetics at submaximal Ca^2+^ and/or a change in the force transmission pathway. Taken together, we conclude that MyBP-C^C1C7^ domains play an important role in contractile performance which helps explain why mutations in these domains often lead to debilitating diseases.

---

Muscle contraction is produced via the interaction of myofilaments^1,2^ and is regulated so that muscle performance matches demand requirements^3–5^. Myosin-binding protein C (MyBP-C) is thought to control muscle contraction via regulation of myosin motors and to form a link between the myofilaments (“C-links”)^3,6–8^. Here we controllably remove a portion of fast-isoform MyBP-C *in-situ* and provide mechanical and structural evidence demonstrating the impact of MyBP-C on myosin function^6,9^, and the existence of C-links. Mutations of MyBP-C are long known to be involved in the etiology of debilitating skeletal and cardiac muscle diseases^9–11^, which our study helps detail to the benefit of future treatment strategies.

Muscle sarcomeres are composed of an interdigitating array of myosin-containing thick and actin-containing thin filaments (Fig. 1A). Contraction force is generated via a carefully orchestrated action where myosin-based motor proteins of the thick filaments interact with the actin molecules of the thin filaments to generate force and length changes by a process called crossbridge cycling^1,2^. Myosin molecules are hetero-multimers and have a tail, neck (regulatory and essential light chain region) and head (motor) regions, which are packed into the thick filament as a three stranded quasi-helical array forming repeating “crowns” of three sets of myosin heads separated by ∼120° from each other^12^. As one moves along the thick filament axially, crowns arise at ∼14.3 nm intervals, each rotated by 40°, so that the crown orientation repeats every three turns / ∼43 nm (Fig. 1B). Within this repeat, each of the three myosin crowns has slightly different features and are denoted here as Cr_1_, Cr_2_, and Cr_3._ Thick filament crowns are also classified as occurring in 3 distinct segments of the thick filament, with 6 crowns in the P-(proximal), 27 crowns in the C-(central), and 18 crowns in the D-(distal) zone, where C-zone crowns associate with an extra protein called myosin-binding protein C (MyBP-C) with a stoichiometry of ∼ three MyBP-C molecules spaced at 43 nm intervals (Fig. 1A-B)^3,13–15^.

**Fig. 1.**
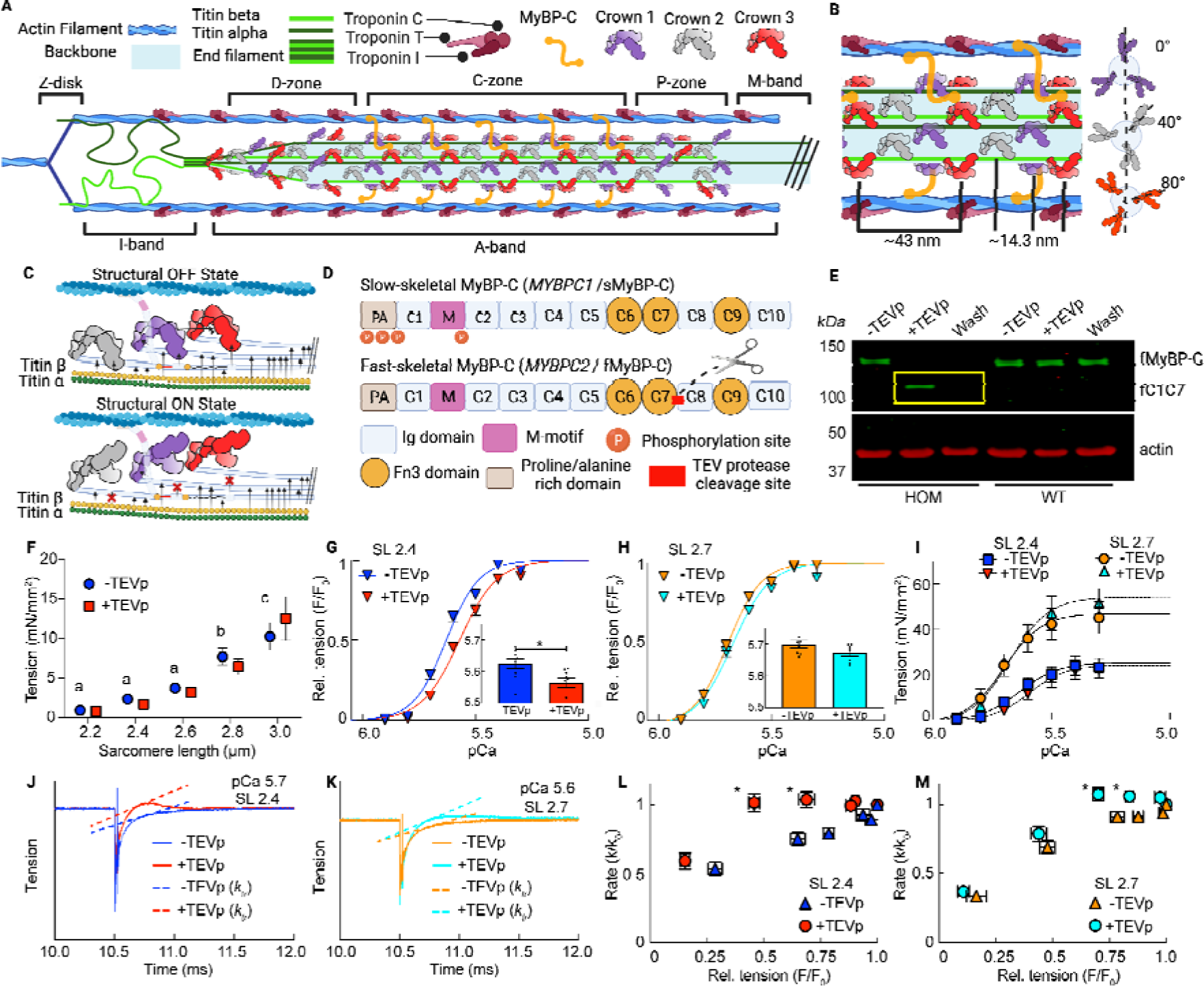
Study of MyBP-C in skeletal muscle by targeted, acute and specific cleavage in situ. (**A**) Schematic of a skeletal half-sarcomere. (**B**) Expanded view of crown placement on thick filaments. (**C**) Schematic of one ∼43 nm repeat in the C-zone, with crowns shown in the structural OFF and ON states (details in text). Arrows indicate the suggested interactions between different elements^39,51,52^. (**D**) Domain layout of fast and slow twitch MyBP-C, with the TEV protease cleavage site of SNOOPC2 mouse line indicated. (**E**) Western blot of tagged fast MyBP-C isoform from homozygous and wildtype SNOOPC2 psoas, before (fMyBP-C) and after fast isoform cleavage (fC1C7 – cleaved N-terminal domains). (**F-M**) Mechanical measurements of permeabilized psoas fibers from SNOOPC2 muscle, before and after TEV protease treatment (N-terminal domains removed) for: passive tension-SL (F), relative tension-pCa curves at 2.4 (G) and 2.7 (H) µm SL, absolute tension-pCa curves (I), representative tension k_tr_ curves before (J) and after (K) TEV protease treatment, and normalized k_tr_ vs. relative tension at 2.4 (L) and 2.7 (M) µm SL. Statistics throughout are repeated-measures ANOVA designs followed by a Tukey Honestly Significant Difference (HSD) post-hoc test on significant main effects, or paired t-tests. *P < 0.05 after treatment at each SL. Letter in (F) indicate differences between SL (no treatment effects to report). Data throughout reported as mean ± s.e.m. Further statistical details in Extended Data Table 1-4.

In addition to the text-book description of a thin filament-based regulation scheme of muscle contraction^16,17^, there is a growing appreciation of thick filament-based regulation mechanisms acting in parallel^5,18^. Structurally, a proportion of myosin heads are in an “OFF” conformation, unable to interact with actin and docked close to the thick filament backbone in quasi-helical tracts. Other myosin heads are in an “ON” state positioned away from the thick filament backbone where they can readily bind to the thin filaments during contraction (Fig. 1C)^5,19^. In resting muscle, most heads are in the OFF state with only a few in the ON state. The strain-dependent thick filament activation model^20^ posits that these few ON state heads generate strain in the thick filament early in activation that causes a cooperative OFF-to-ON transition of myosin heads to participate in contraction. Thick filament activation mechanisms are also likely to be involved in myofilament length dependent activation (LDA), the phenomenon whereby the sarcomeres can generate more force at longer sarcomere lengths (SL) at a given level of activating calcium^21^. LDA is now understood as a general feature of striated muscles and often found to be dysregulated in skeletal- and cardiomyopathies^5,21,22^.

LDA is thought to involve two major players: the large titin proteins that run along the half-thick filament length in the A-band and bridge the thin and thick filaments in the I-band of the sarcomere^23,24^, and the thick filament-bound MyBP-C^3^. However, these molecules could also be involved more generally in mechanisms of thick-filament activation, where the activation sensitivity of crossbridges is tunable^3,23,25^. MyBP-C has three paralogs that are encoded by distinct genes that are generally expressed by muscle type: cardiac (encoded by *MYBPC3*), slow skeletal (sMyBP-C, encoded by *MYBPC1*), and fast skeletal (fMyBP-C, encoded by *MYBPC2*) (Fig. 1D). Mutations in the skeletal muscle paralogs lead to debilitating myopathies in humans including severe and lethal forms of distal arthrogryposis myopathy^10,11^ and lethal congenital contracture syndrome^9^, suggesting a critical role in healthy sarcomere function. In addition, fMyBP-C knockout mice present altered force generation with myosin heads (i.e., free-heads) shifted towards the ON state^9,26^, suggesting that MyBP-C helps stabilize the myosin OFF state and/or suppresses OFF-to-ON transitions^26–28^. Importantly, MyBP-C effects can be blunted via post-translational modifications such as phosphorylation^29–31^, which can be altered in some myopathies but also in healthy muscle during exercise, e.g., via increased PKA/PKC activity^32^. The mechanism(s) by which MyBP-C affects the myosin ON/OFF state are not clear, but the phenomenon is remarkable because there are relatively few MyBP-C (up to ∼54) molecules to regulate the behavior of up to ∼306 myosin heads per thick filament.

To evaluate the structural and functional effects of MyBP-C in sarcomeres, we developed the SNOOPC2 mouse line (Fig. 1D, Extended Data Fig. 1A), which has an engineered tobacco etch virus protease (TEV_P_) recognition site inserted between the C7 and C8 domains of the *MYBPC2* gene. This addition allows for immediate, controllable, and specific cleavage of the N-terminal region of fMyBP-C (from the N-terminal proline/alaline sequence through the C7 domain; MyBP-C^C1C7^) in permeabilized skeletal muscle, leaving only the C-terminal portion (C8-C10) anchored to the thick filament. This powerful experimental approach allows for the study of changes that occur before and after removal of MyBP-C^C1C7^ *within the same preparation*, removing variation caused by using different samples between conditions and improving statistical power. Here we used psoas muscle from homozygous SNOOPC2 mice for this evaluation because nearly all fibers are of fast-twitch composition^33^. Western blots confirmed successful cleavage of fMyBP-C in permeabilized homozygous SNOOPC2 psoas (Fig. 1E), while not targeting sMyBP-C (Extended Data Fig. 1B) or wildtype fMyBP-C (Fig. 1E). Furthermore, passive tension of fMyBP-C was not affected by TEV_P_ treatment in fibers from homozygous or wildtype mice in relative or absolute isometric tension (Fig. 1F; Extended Data Fig. 1C).

We evaluated the tension-pCa (-log[Ca ^2+^]) relationship in permeabilized fiber bundles before and after cleavage of N-terminal domains of fMyBP-C at shorter (2.4 µm) and longer (2.7 µm) SL (Fig. 1G-I; Extended Data Fig. 1D). Cleavage caused a significant rightward shift of the tension-pCa relationship, as quantified by the pCa_50_ (i.e., the pCa at which active force was half-maximal), at shorter but not at longer SL, indicating a decreased calcium sensitivity at shorter lengths (Fig. 1G-H). Although pCa_50_ was altered, LDA was observed from short to long SLs (i.e., increasing pCa_50_) in samples both before and after MyBP-C^C1C7^ removal. While cleavage significantly affected tension at submaximal activation levels (pCa > 5) at shorter SL, supramaximal activation (pCa 4.5) tension was affected at longer SL (Fig. 1I). Next, we assessed the rates of force redevelopment following a slack/restretch maneuver (*k*_tr_), a measure of crossbridge cycling, across a range of calcium concentrations (Fig. 1J-K). At 2.4 and 2.7 µm SL, the *k*_tr_ vs. force relationships were similar to previous studies^27,34^, with *k*_tr_ increasing with relative force (Fig. 1I-J). Qualitatively, cleaved samples produced a consistent overshoot of the tension before settling to expected values (Fig. 1I-J), which has been reported previously^35^. At both SLs the loss of fMyBP-C^C1C7^ significantly increased *k*_tr_ at intermediate but not low or maximal active force levels (Fig. 1L-M). Furthermore, as with calcium sensitivity, *k*_tr_ was less affected by fMyBP-C cleavage at longer vs. shorter SLs. These data are indicative of faster rates of myosin crossbridge cycling after cleavage of fMyBP-C at intermediate calcium activation, consistent with effects observed following loss of cardiac MyBP-C^C1C7^ in cardiomyocytes, and in cardiac and fast skeletal knockout mice^27,34^. Detailed statistical information for mechanical datasets is provided in Extended Data Tables 1-4. Taken together, these data show that fMyBP-C^C1C7^ modulate force generation and crossbridge cycling kinetics, specifically at moderate levels of activation (∼40-80% of maximal force), in agreement with previous observations in a cardiac MyBP-C TEVp cleavage model, knockout mouse models, and *in vitro* biochemical studies_3,9,10,27_.

To determine structural effects in sarcomere proteins due to fMyBP-C^C1C7^ removal, we used small-angle X-ray diffraction, a powerful method that leverages the partially crystalline arrangements of sarcomere proteins to evaluate their structural features under near-physiological conditions^5,36^. A focused, high intensity X-ray beam from a synchrotron source passes through a fiber perpendicular to its long axis, producing a diffraction pattern on a detector. The intensities and spacings of the diffraction features (i.e., reflections; Fig. 2A) provide structural information regarding specific periodic features in the sarcomere, as described below. The equatorial portion of the X-ray diffraction pattern provides information pertaining to the sarcomere lattice, while the meridional portion provides information regarding axial periodicities in the protein arrangement in the thick and thin filaments^5,37,38^. Control experiments with wildtype preparations incubated with TEV_P_ indicated no TEV_P_ impact on structural features (Extended Data Fig. 2) and detailed statistical information for control and experimental datasets are provided in Extended Data Table 5-7.

**Fig. 2.**
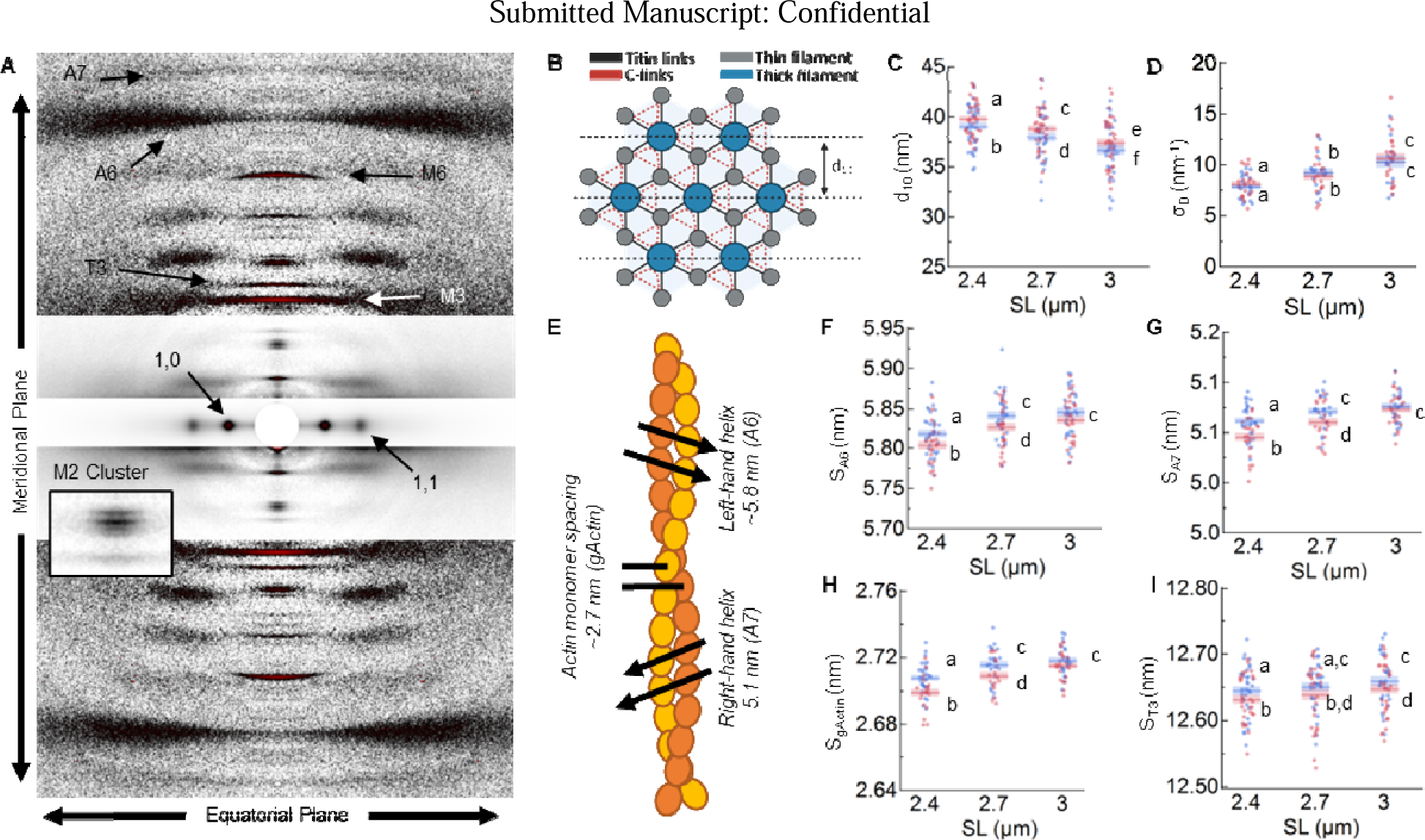
Lattice and thin filament structural parameters of skeletal muscle fibers before (blue) and after (red) cleavage of the fMyBP-C bridge region across SLs. (A) A representative X-ray diffraction pattern of permeabilized fiber bundles, with key reflections and axis orientation indicated. Three different intensity scales were overlayed so that features of interest could be best viewed by eye. (B) Schematic of the sarcomere lattice, with titin and MyBP-C C-link points indicated, as well as the d_10_ lattice plane. C-links can attach to either of the 2 actin filaments nearby, and so this is depicted by the orange triangle. (C) Quantified d_10_. (D) Quantified σ_D_, a measure of the variability in the d_10_ spacing. (E) Cartoon representation of actin, with important structural features indicated. (F) A6 spacing (S_A6_) from the right-handed actin helix. (G) A7 spacing (S_A7_) from the left-handed helix of actin. (H) Actin monomer spacing (S_gActin_), a measure of thin filament axial length. (I) T3 spacing (S_T3_) from the troponin periodicity. Statistics throughout are ANOVA designs with main effects treatment, SL, and their interaction, and a random effect of individual, followed by Tukey’s Honestly Significant Difference (HSD) post-hoc test on significant main effects (P < 0.05), and reported in figures as connecting letters: different letters are significantly different. Data throughout reported as mean ± s.e.m. Further statistical details in Extended Data Table 5.

We first considered a long-debated question in muscle physiology: are there load bearing MyBP-C “C-links” between thick and thin filaments?^13,39,40^ The interdigitating thick and thin filaments of the sarcomere form a hexagonal lattice, where 6 thin filaments surround a thick filament (Fig. 2B). Mechanically speaking, in passive muscle, any protein linking the thick to thin filaments should pull on the thin filaments as SL increases, which would be expected to produce both radial forces that pull the filaments together and longitudinal forces that elongate the thin filaments, both measurable in our X-ray experiments. The inter-filament lattice spacing in relaxed sarcomeres is a key determinant of force production because crossbridge kinetics are sensitive to lattice spacing^41,42^. Lattice spacing is well known to be modulated by titin filaments, specifically the I-band spring region that extends from the actin filaments near the Z-line to the ends of thick filaments. I-band titin-based tension increases with increasing SL, producing a compressive force on the myofilament lattice and centering the thick filaments within the sarcomere^23,43–45^. Inter-filament lattice spacing can be derived from the 1,0 equatorial reflections, while the radial width of the peaks estimated via the radial width parameter, σ_D_, provides a measure of lattice spacing inhomogeneity^5^ among myofibrils. Before treatment, increasing SL increased σ_D_ (Fig. 2D) and decreased d_10_ (Fig. 2C) as previously reported^23,43^. In comparison, fMyBP-C^C1C7^ removal did not alter σ_D_ (Fig. 2D) but d_10_ generally increased across SLs (Fig. 2C) suggesting that fMyBP-C exerts a radial force opposing the expansion of the lattice, in agreement with a previous report on fMyBP-C KO muscle^27^. Our data provide compelling complementary evidence that MyBP-C does indeed form C-links with the thin filament^13,39,40^.

We next turned our attention to SL-dependent thin filament elongation, which has previously been observed in passively stretched cardiac and skeletal sarcomeres^23,46^. Here, we measured thin filament elongation (Fig. 3E) using the spacing of the left-handed helix, S_A6_; at ∼5.8 nm; (Fig. 2F) and the right-handed helix, S_A7_; at ∼5.1 nm; (Fig 2G) in the actin filaments to estimate (see Methods) the axial spacing of actin monomers, S_gActin,_ at ∼2.7 nm (Fig. 2H). Before treatment, S_gActin_ increased with SL as described previously^23^, with thin filament extension observed from 2.4 to 2.7 µm SL with no further elongation from 2.7 to 3.0 µm SL (Fig. 2I). In comparison, fMyBP-C^C1C7^ removal significantly reduced S_gActin_ at 2.4 and 2.7 µm SL, but not at 3.0 µm SL (Fig. 2H). However, these changes did alter the magnitude of thin filament elongation from 2.4 to 2.7 µm SL, while elongation also continued from 2.7 to 3.0 µm SL to match the 3.0 µm SL values before treatment. We additionally assessed the spacing of the third order meridional reflection (S_T3_; Fig. 2I) from the thin filament bound troponin complex. S_T3_ also increased with thin filament elongation following fMyBP-C^C1C7^ removal which could be completely accounted for by the stretching of the thin filament (ANCOVA covariate P < 0.0001, treatment P = 0.28; interaction P = 0.93) indicating no detectable structural changes in troponin apart from that imposed by thin filament stretch itself.

**Figure 3.**
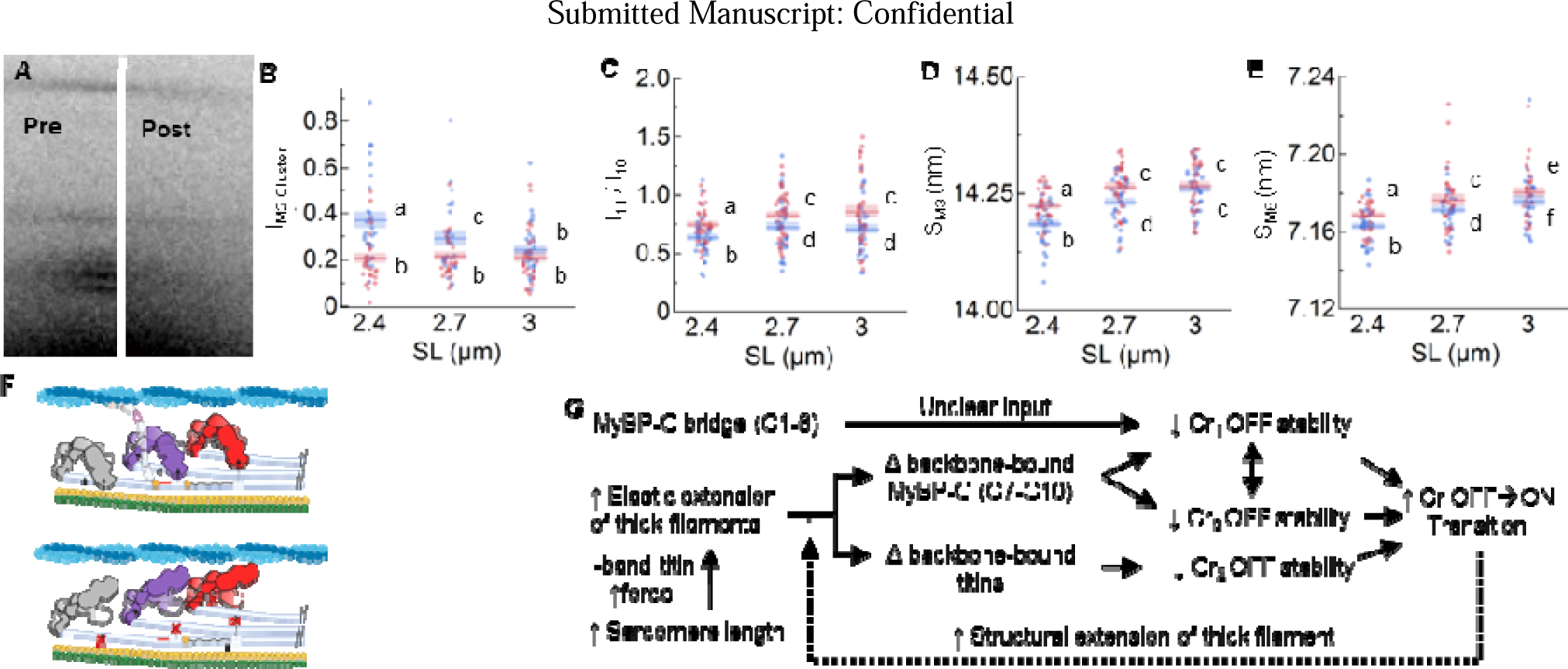
Myosin ON/OFF structural parameters of skeletal muscle fibers before (blue) and after (red) cleavage of the fMyBP-C bridge region across SLs. (**A**) M2 cluster before and after fMyBP-C cleavage. (**B**) Intensity of the M2 cluster (I_M2_) cluster. (**C**) I_1,1_/I_1,0_ provides a measure of the mass distribution between the thick and thin filaments. (**D**) M3 spacing (S_M3_) measures the orientation of myosin heads. (**E**) M6 spacing (S_M6_) is a measure of thick filament length. (**F**) Representation of myosin OFF-ON transition before (top) after (bottom) N-terminal fMyBP-C cleavage. We predict that a loss of regulation of N-terminal MyBP-C leads to an OFF→ON transition on Cr_1_ (purple) myosins has a cooperative effect on surrounding Cr_2_ and Cr_3_ myosins to also transition OFF→ON. (**G**) Based on mechanical and structural details from the current and other studies, we present a flow chart of myosin head ON/OFF control. This chart shows how myosin heads transition OFF→ON, but we assume the reverse effects will also transition ON→OFF. Statistics throughout are ANOVA designs with main effects treatment, SL, and their interaction, and a random effect of individual, followed by Tukey’s Honestly Significant Difference (HSD) post-hoc test on significant main effects (P < 0.05), and reported in figures as connecting letters: different letters are significantly different. Data throughout reported as mean ± s.e.m. Further statistical details in Extended Data Table 6.

It is remarkable that while fMyBP-C^C1C7^ removal decreased overall thin filament length, it did not eliminate thin filament elongation during passive sarcomere stretch. We posit three possibilities that are not necessarily mutually exclusive. First, the uncleavable slow-MyBP-C^C1C7^s, although few in psoas muscle, could nevertheless form C-links and contribute to stretch-dependent thin filament extension. The second relies on the property that low-level crossbridge formation occurs in passive muscle and so can contribute to thin filament stretch. It is possible that fMyBP-C^C1C7^ removal increases this behavior, as is supported by our structural data below. Third, because lattice spacing increases after fMyBP-C^C1C7^ removal, titin filaments are stretched thereby increasing titin-based forces on the thin filament, at least where titin interacts with the thin filament in the I-band. The potential interplay between these scenarios makes it difficult to study each individually but could potentially be separated out using computational modeling and are worth exploring^47^.

Of note for this study is a grouping of up to four X-ray meridional reflections related to the presence of MyBP-C in the so-called M2 cluster^38,48^ that are not present in computed transforms of electron micrographs, analogous to X-ray diffraction patterns from MyBP-C KOs^7^. In the present study, we did not typically have sufficient resolution to separate out the 4 reflections and so instead we measured the intensity of this entire M2 cluster^49^ (I_M2 cluster_; Fig. 3A) before and after removal of fMyBP-C^C1C7^. We report that prior to treatment, increasing SL decreased I_M2 cluster_. In contrast, fMyBP-C^C1C7^ removal substantially decreased I_M2 cluster_ at shorter SLs to a value that was independent of SL (Fig. 3A-B). While the M2 cluster is clearly associated with the presence of fMyBP-C^C1C7^, the reflections are most likely not due to MyBP-C molecules themselves because MyBP-Cs are relatively sparse compared to other repeating proteins (i.e., myosins) and so would be expected to create a reflection too weak to be measured^37^ Instead, the M2 cluster reflections most likely arise from a sub-population of myosin heads that interact with MyBP-C in the C-zone, which causes them to be perturbed and generate their own periodicities (e.g. the M2 cluster)^37,50^. In complementary evidence, high-resolution cryo-EM and cryo-ET structures of cardiac thick filaments in the OFF state indicate interaction and potential cooperativity of myosin heads with other myosin heads and MyBP-C^39,51,52^, providing a pathway for relatively few MyBP-Cs to impact many myosin head ON/OFF states. In summary, our data supports the notion that the fMyBP-C^C1C7^ contributes to the M2 cluster reflections by binding and perturbing a subpopulation of myosin heads.

There are structural signatures that provide evidence for MyBP-C stabilizing the OFF conformation of the myosin heads in the C-zone^3,27,28^ as previously proposed^39^. The equatorial intensity ratio (I_1,1_/I_1,0_) is often used to quantify the myosin head OFF/ON state transitions. It provides a measure of the transfer of mass (i.e., myosin heads) from the thick to the thin filaments, with increasing I_1,1_/I_1,0_ indicating that more myosin heads are associated with the thin filaments^53^. In additional complementary evidence, myosin head configuration can be evaluated via the spacing of the M3 myosin meridional reflection (S_M3_), which reflects the average distance of ∼14.3 nm spacing between the myosin crowns along the thick filament. Increases in S_M3_ are typically associated with a subpopulation of myosin heads moving from the OFF to the ON state or the reorientation of myosin heads to a more perpendicular orientation relative to the thick filament backbone that increases the chance of attachment^5,54^. We report that, before fMyBP-C^C1C7^ removal, I_1,1_/I_1,0_ (Fig. 3C) and S_M3_ (Fig. 3D) behaved as expected^23^ by shifting to larger values when fibers were stretched from 2.4 to 2.7 µm SL, indicating that a proportion of myosin heads shifted from the OFF to the ON state in response to passive stretch. After N-terminal fMyBP-C cleavage, both I_1,1_/I_1,0_ and S_M3_ were significantly elevated at 2.4 and 2.7 µm SL but still followed the same length-dependent trend as before cleavage. Since the structural and mechanical signatures (Fig. 1I) of LDA are still present after fMyBP-C cleavage, we conclude that other mechanisms dominate in LDA such as those involving the fMyBP-C C-terminal domains (C8-C10)^39^ and strain generated in the thick filaments by titin-based passive tension^23,43,46^.

Compared to the length-dependent increase of I_1,1_/I_1,0_ and S_M3_ from 2.4 to 2.7 µm SL, there is no detectable increase from 2.7 to 3.0 µm SL (Fig. 3C-D). However, this may not suggest that the LDA effect has reached a maximum. The decrease in thick-thin filament overlap associated with increasing the SL from 2.7 to 3.0 µm SL can itself decrease I_1,1_/I_1,0_, regardless of myosin head movement^55^, while myosin heads in the ON state potentially become disordered and may contribute less to S_M3_ compared to OFF myosin, pushing values down. Therefore, the probable structural scenario represented by the data is that more myosin heads are moving radially towards the thin filament with increasing SL between 2.4 and 2.7, as well as from 2.7 and 3.0 µm SL (OFF-to-ON transitions). Taken together, our data support the notion that loss of fMyBP-C^C1C7^ prevents it from stabilizing the OFF state of myosin heads, resulting in release of some myosin heads from the OFF to the ON state. This would also explain the results of MyBP-C phosphorylation, which also leads to OFF-to-ON transitions of myosin heads^28,56^ and may be caused by a phosphorylation dependent destabilization of MyBP-C and myosin heads.

We next focused on the elongation of thick filaments associated with sarcomere-stretch in passive muscle, a property correlated with increasing OFF-to-ON myosin heads transitions that has been proposed to contribute to LDA in some systems^5,19,23^. Elongation of the thick filaments during relaxed sarcomere stretch is predominately caused by titin-based forces, which pull on the tips of the thick filaments and increase with increasing SL^23,43^. Thick filament extension can be quantified by the spacing of the M6 reflection (S_M6_), which tracks the ∼7.2 nm periodicity that arises off the thick filament backbone, with increasing S_M6_ indicative of thick filament extension^57,58^. Before and after fMyBP-C^C1C7^ loss, we observed the expected increase in S_M6_ with stretch from short to long SLs (Fig 3E). Following cleavage of fMyBP-C, however, S_M6_ was increased at every SL, suggesting that fMyBP-C^C1C7^ affects thick filament structure independent of the titin-based forces placed on it. Of note, the regression slope of S_M6_ versus SL is similar between treatment conditions (treatment*SL P = 0.62), suggesting that changes in filament stiffness do not play a role in thick filament structural changes in this case.

This begs the question: what drives this filament stiffness- and SL-independent change in thick filament length? Recent studies provide evidence that simply shifting a subset of myosin heads between OFF and ON states via chemical treatment of relaxed fibers can change thick filament length^4,19,38,59^. For example, incubation with the drug mavacamten transitions myosin heads from the ON to the OFF state and is accompanied by a shorter S_M6_^19^, while incubation with 2’-deoxy-ATP (dATP) transitions myosin heads from the OFF to the -ON state and is accompanied by a longer S_M6_^4^. Our data indicate that loss of fMyBP-C^C1C7^ promotes OFF-to-ON transitions (increases in I_11_/I_10_ and S_M3_), and so this would also track with changes to thick filament length. The idea that MyBP-C can regulate the OFF-to-ON transition of myosin heads has been discussed for some time^5,20,60^ but our data (Fig. 3C-F) combined with recent sarcomere and thick filament reconstructions^39,52^ provide sufficient detail to suggest a possible mechanism (Fig. 3G). As detailed in Figure 3G, our data are consistent with the hypothesis that loss fMyBP-C^C1C7^ promotes OFF-to-ON transitions of Cr_1_ myosin heads in the C-zone. This is proposed to create localized elongations of the thick filament backbone (which we observe) that affect nearby OFF Cr_2_ and Cr_3_ myosin heads, which then break their docked formations and transition from OFF-to-ON states. One critical detail to uncover is why this purported feed-forward mechanism does not continue until all myosin heads are in the ON state, implying the existence of some type of as-of-yet unknown molecular brake to limit OFF-to-ON state changes.

Combining our data with OFF-to-ON transition data in titin^23^, myofilament protein location, and interaction data from recent high resolution cryo-ET experiments^39,51,52^, and 30 years of accumulated data and hypotheses regarding the functional roles of MyBP-C^3^, we propose a unifying scheme for sarcomere OFF-to-ON transitions (Fig. 3F-G). Sarcomere stretch produces titin-based forces that elongate the thick filament backbone, leading to strain in the thick filament-bound titin and MyBP-C domains. This disrupts the myosin OFF-state, namely those involving Cr_2_ (titin interactions^39^ across the whole filament, and Cr_1_ and Cr_3_ interactions in the C-zone (MyBP-C C8 and C10, respectively^39,52^). Separately, N-terminal MyBP-C domains act to maintain myosin heads in the OFF state, but when perturbed (in our case, removed), there are OFF-to-ON transitions of Cr_1_ myosin heads, which are thought to then indirectly destabilize Cr_3_ myosin OFF states via the stabilizing interactions of Cr_1_ or Cr_3_ myosins^51^. Furthermore, OFF-to-ON transitions of Cr_1_ lead to a sarcomere-length independent structural elongation of the thick filament^4,19,20^, which perturbs thick-filament bound titin and MyBP-C domains, potentially destabilizing their interaction with docked myosin heads. It should also be noted that myosin, MyBP-C, and titin can all be affected in ways that impact their role on OFF-ON myosin states by mutations or by local environmental factors such as phosphorylation, oxidation, and pH — all of which can occur in disease but also are common in healthy people during exercise^61–63^. Therefore, there may be many ways to fine tune OFF-to-ON state transitions in skeletal muscle to meet performance demands in real-time and provide plasticity in health and disease.

## Acknowledgments

We thank the BioCAT beamline support staff at the APS, Massimo Reconditi, and Anna Good. BioRender.com was used to construct some of the figure cartoons. This research used resources of the Advanced Photon Source, a U.S. Department of Energy (DOE) Office of Science User Facility operated for the DOE Office of Science by Argonne National Laboratory under Contract No. DE - AC02 - 06CH11357, and further NIH support. The content is solely the responsibility of the authors and does not necessarily reflect the official views of the National Institute of General Medical Sciences or the National Institutes of Health. Funding for this study was provided by the German Research Foundation (454867250 [ALH], SFB1002A08 [WAL]), IZKF Münster (Li1/029/20 [WAL]), National Institute of Health (P41 GM103622 [TCI]), P30 GM138395 [TCI], HL080367 [SPH]), HL140925 [SPH], AR081935 [SPH], T32 [NME], American Heart Association (827628 [NME]).

## Author Contributions

A.L.H AND S.P.H. Conceptualized the project; A.L.H., W.M., N.M.E., R.S., S.P.H. developed the methods; A.L.H., N.M.E., R.S., D.N., M.K., S.P.H. conducted the investigation; A.L.H., N.M.E., M.K. visualized the datasets; A.L.H., S.P.H., T.C.I., W.A.L.; A.L.H. was the project administrator; A.L.H., S.P.H. supervised the study; A.L.H. wrote the original draft; all authors review, edited, and agreed to the final draft.

## Additional Information

Supplementary information is available for this paper. Correspondence and requests for materials should be addressed to A.L.H. or S.P.H. Reprints and permissions information is available at www.nature.com/reprints.

## Data and materials availability

All data are available in the main text or the supplementary materials, or available upon reasonable request.

## Methods

### Animal model and muscle preparation

#### SNOOPC2 mice

Animal procedures were approved and performed according to the guidelines of the local animal care and use committee (IACUC) of the University of Arizona. SNOOPC2 mice were bred and housed at the University of Arizona. Genotyping was performed via PCR analysis and protein gels used to assess TEV protease reactivity evaluated by measuring myosin binding protein C (MyBP-C) cleavage before and after treatment. Genetically homozygous and wildtype adult SNOOPC2 mice (age range, 2 – 6 months) were humanely euthanized; psoas muscle was immediately extracted for long-term storage and permeabilized (“skinned”) at −20°C using standard glycerol techniques (1:1 rigor: glycerol; rigor contains (in mM) KCl (100), MgCl_2_ (2), ethyleneglycol-bis(β-aminoethyl)-N,N,N, N-tetraacetic acid (EGTA,5), Tris (10), dithiothreitol (DTT, 1), protease inhibitors [Complete, Roche Diagnostics, Mannheim, Germany], pH 7.0). Samples were shipped to the BioCAT facility on ice for all experimental tests and stored at −20°C until used. On the day of experiments, psoas muscles were removed from the storage solution and vigorously washed in relaxing solution (composition (in mM): potassium propionate (45.3), N,N-Bis(2-hydroxyethyl)-2-aminoethanesulfonic acid BES (40); EGTA (10), MgCl_2_ (6.3), Na-ATP (6.1), DTT (10), protease inhibitors [Complete], pH 7.0)). Bundles containing 10-20 fibers (3-6 mm long) were carefully excised and kept in physiological register by tying silk suture knots (sizing 6-0 or 4-0) at the distal and proximal ends of the bundle. Samples were then immediately transferred to the experimental chamber (see below).

### Experimental protocols and analysis

#### N-terminal fMyBP-C cleavage

All experiments were conducted by running the below mechanical experiments before and after incubation with tobacco etch virus (TEV) protease, with selectively cleaves the TEV protease recognition site of fast-isoform MyBP-C in SNOOPC2 muscle so that the N-terminal region detaches and diffuses out of the sarcomere (Fig. 1; Extended Fig. 1), similar to that designed previously for cardiac MyBP-C^1^. The samples were incubated with TEV_P_ for 30 mins. Recombinant TEVp was purified as previously described^1^ or was purchased from Thermo Fisher Scientific, USA and used at 100 units acTEV_P_ in 300 μl relaxing solution. After incubation, fibers were rinsed in fresh relaxing solution to remove excess protease.

#### Western Blot

Proteins were prepared for western blotting by pulverizing left ventricle tissue from both WT and HOM SnoopC2 mice with a pestle and mortar cooled with liquid nitrogen. Left ventricle tissue was then homogenized using a Polytron Homogenizer (PT1200E, Kinematica, Switzerland) in a skinning solution ([in mmol/L]: 5.92 Na_2_ATP, 6.04 MgCl_2_, 2 EGTA, 139.6 KCl, 10 imidazole with 0.01% saponin, 1% Triton X-100®, and Halt protease inhibitor cocktail, EDTA-free [78437, ThermoFisher Scientific, USA], pH 7.0). The homogenized tissue was then tumbled for 15 minutes at 4°C and washed in relaxing solution ([in mmol/L]: 5.92 Na_2_ATP, 6.04 MgCl_2_, 2 EGTA, 139.6 KCl, 10 imidazole). Tissue was then either left untreated, treated with TEV protease (12 μg protease per mg tissue) for 30 minutes at room temperature, or treated with TEV protease for 30 minutes at room temperature and then washed in relaxing solution to remove cleaved protein. The tissue in relaxing solution was then mixed with an equal volume of urea buffer ([in mol/L]: 8 urea, 2 thiourea, 0.05 Tris-HCl, 0.075 dithiothreitol with 3% SDS and 0.03% bromophenol blue, pH 6.8), run on an SDS-PAGE gel (4561086, 4-15% Mini-PROTEAN® TGX^TM^ Precast Protein Gel, Bio-Rad), and transferred onto a nitrocellulose membrane. Blots were blocked with OneBlock^TM^ Fluorescent Blocking Buffer (20-314, Genessee Scientific) and stained for either fMyBP-C (MYBPC2 polyclonal rabbit antibody diluted 1:2000, PA5-83638, ThermoFisher Scientific, USA) or sMyBP-C (MYBPC1 polyclonal rabbit antibody diluted 1:1000, NBP2-41157, Novus Biologicals) with actin (actin monoclonal mouse antibody diluted 1:2000, ACTN05 [C4] MA5-11869, ThermoFisher Scientific, USA) as a loading control. Secondary antibodies used were goat anti-rabbit IRDye 800CW (926-32211, LI-COR) and goat anti-mouse IRDye 680RD (926-68070, LI-COR).

#### Crossbridge kinetics/mechanics

For mechanical measurements in permeabilized psoas from SNOOPC2 mice, psoas myocytes were mechanically homogenized using a Polytron Homogenizer (PT1200E, Kinematica, Switzerland) in a skinning solution (5.92 mM Na_2_ATP, 6.04 mM MgCl_2_, 2 mM EGTA, 139.6 mM KCl, 10 mM imidazole, 0.01% saponin, 1% Triton-X-100®, Halt protease inhibitor cocktail, EDTA-free [78437, Thermo Fisher Scientific, USA]). Following mechanical homogenization, cells tumbled for 40 minutes at 4°C to allow for removal of sarcoplasmic reticulum, sarcolemma, and any remaining endogenous Ca^2+^, leaving intact the myofibrillar network. Psoas cells were then washed to remove remaining detergents of the skinning solution. Psoas cells (∼100-250 µm in length) were attached between a high-speed motor (Model: 315C-I, Aurora Scientific Inc., Aurora, Ontario, Canada) and a force transducer (Model 403A series, Aurora Scientific Inc., Aurora, Ontario, Canada) with an aquarium sealant (Marineland, 100% clear silicone rubber). The motor and force transducer were mounted above a temperature-controlled platform (Model 803B, Aurora Scientific Inc., Aurora, Ontario, Canada) regulated by a thermocouple (825A, Aurora Scientific Inc., Aurora, Ontario, Canada), located on the stage of an inverted microscope (Model IX-53, Olympus Instrument Co., Japan). The glue was allowed to cure for 30 minutes before experiments were started. Using push-button micromanipulators, sarcomere length (SL) was set to either 2.4 or 2.7 µm determined from video analysis and force measurements were made by activating myocytes in pCa solutions containing variable free calcium concentrations, ranging from pCa 9.0 to 4.5. All measurements were taken at 15°C. Isometric force measurements (F) were normalized to maximal force (F_0_) at pCa 4.5 and to cross-sectional area of the muscle preparation assuming circular dimensions. Data was plotted in GraphPad Prism and fitted using a sigmoidal 4 parameter logistic curve, as described previously ^1^. The pCa_50_ is the concentration of Ca^2+^ required to achieve half-maximal activation of the myocyte. The rate of force redevelopment (k_tr_) was calculated by fitting force traces with a single exponential curve and normalizing the rates (k) to the maximal rate of force redevelopment (k_0_) measured at pCa 4.5. All rates were then plotted against isometric force. Passive force values were measured in a range of SL (2.2, 2.4, 2.6, 2.8, and 3.0 µm) and plotted as absolute values or normalized to the maximal passive force measured at SL 3.0 µm.

#### Small-angle X-ray diffraction experiments

X-ray diffraction patterns were collected using the small-angle instrument on the BioCAT beamline 18ID at the Advanced Photon Source, Argonne National Laboratory^2^. The X-ray beam (0.103 nm wavelength) was focused to ∼0.06 x 0.15 mm at the detector plane, with an incident flux of ∼3×10^12^ photons per second. The sample to detector distance was set at ∼2 m, and the X-ray fiber diffraction patterns were collected with a downstream CCD-based X-ray detector (Mar 165, Rayonix Inc, USA). Muscle preps were hung on custom muscle mechanics rigs, as explained previously^3^. For TC experiments, diffraction patterns were captured with 1 s exposure times. An inline camera built into the system allowed for initial alignment with the X-ray beam and continuous sample visualization during the experiment. SL (SL) was measured via laser diffraction using a 4-mW Helium-Neon laser. Force baseline was set at slack length. After this initial setup, fiber length changes were accomplished through computer control of the motor. Experiments were conducted at 25°C. The mechanical rig was supported on a custom designed motorized platform that allowed placement of muscle into the X-ray flight path and small movements to target X-ray exposure during experiments. Using the inline camera of the X-ray apparatus, the platform was moved to target the beam at different locations along the length of the sample. To limit X-ray exposure of any one part of the preparation, no part of the sample was exposed more than once. Fiber diameter was measured using the inline camera, and physiological cross-sectional area calculated at initial fiber length, with the assumption that the sample was a uniform cylinder longitudinally.

The mechanical protocol during these experiments consisted of passive ramp-hold stretches. Relaxed samples started at 2.4 µm SL, at which an X-ray diffraction pattern was collected, then stretched (over 60 s) to 2.7 µm SL, held for 60 s before another X-ray diffraction pattern collection, and then similarly stretched to 3.0 µm SL and held for a final X-ray diffraction pattern collection. Samples then underwent the TEV protease protocol at 2.4 µm SL as explained above, and the mechanical experiment repeated.

#### X-ray image analysis

X-ray diffraction patterns were initially reduced and prepared for analysis using Bulb (Accelerated Muscle Biotechnologies, Boston, USA), and then analyzed using the MuscleX open-source data reduction and analysis package^4^. The “Scanning Diffraction” routine was used to measure the angular divergence of the 1,0 equatorial reflection. The routine obtains 2D and 1D radially integrated intensities of the equatorial intensities, and then fits Gaussian functions over the diffraction peaks to calculate the standard deviation (width σ) intensity distribution pattern. In this process, the routine obtains the integrated intensity of each equatorial reflection as a function of the integration angle. The “Equator” routine of Muscle X was used to calculate the I_1,1_ / I_1,0_ intensity ratio, lattice spacing (LS) between thick filaments, and σ_D_, a measure of the variability in thick filament lattice spacing (a proxy for lattice ordering). There are limitations to the I_1,1_/I_1,0_ in protocols that disrupted lattice order, such as cleaving titins ^3^ because it is not easy to uncouple the effects of lattice disorder^5,6^ and mass shift due to myosin head movement on I_1,1_/I_1,0_. In the current model, cleaving N-terminal MyBP-C does not show effects on lattice order (e.g. σ_D_) and so we considered I_1,1_/I_1,0_ a trustworthy assessment of the average movement of myosin head mass. Meridional (I_M2_Cluster_, S_M3_, I_M3_, S_T3_, S_M6_) and off-meridional reflections (S_A6_, S_A7_) were collected using the MuscleX routines “Diffraction Centroids” and “Projection Traces”. The S_A6_ and S_A7_ report on the left- and right-handed actin helical structures within the thin filament were used here to calculate the axial spacing of the actin monomers (S_gActin_)^7,8^, where S_gActin_ can be used as a measure of thin filament extension^9,10^. Every image provides reflections of different quality, which lead to various levels of Gaussian fit errors for each reflection modeled, which increases the variation in spacings in the dataset. To limit these effects, fit errors > 10% were discarded. Positions of X-ray reflections on the diffraction patterns in pixels were converted to sample periodicities in nm using the 100-diffraction ring of silver behenate at d_001_ = 5.8380 nm.

### Statistics

Statistical analysis was conducted using JMP Pro (V16, SAS Institute Inc., Cary, NC, USA). Significance level was α = 0.05. Response variables included all X-ray parameters. We first built a repeated-measures analysis of variance (ANOVA) design. We used fixed effects treatment (pre /post TEV protease incubation) and SL, a treatment x sarcomere interaction term, and a random (repeated-measures) effect of individual. Data were best Box-Cox transformed to meet assumptions of normality and homoscedasticity, when necessary, which were assessed by residual analysis, Shapiro-Wilk’s test for normality, and Levene’s test for unequal variance.

Significant main effects were subject to Tukey’s highly significant difference (HSD) multiple comparison procedures to assess differences between factor levels. These data were indicated in graphs via so-called connecting letters, where factor levels sharing a common letter are not significantly different from each other. Data is presented at mean ± standard error of the mean (s.e.m.) unless otherwise noted.

---

**Extended Data Figure 1.**
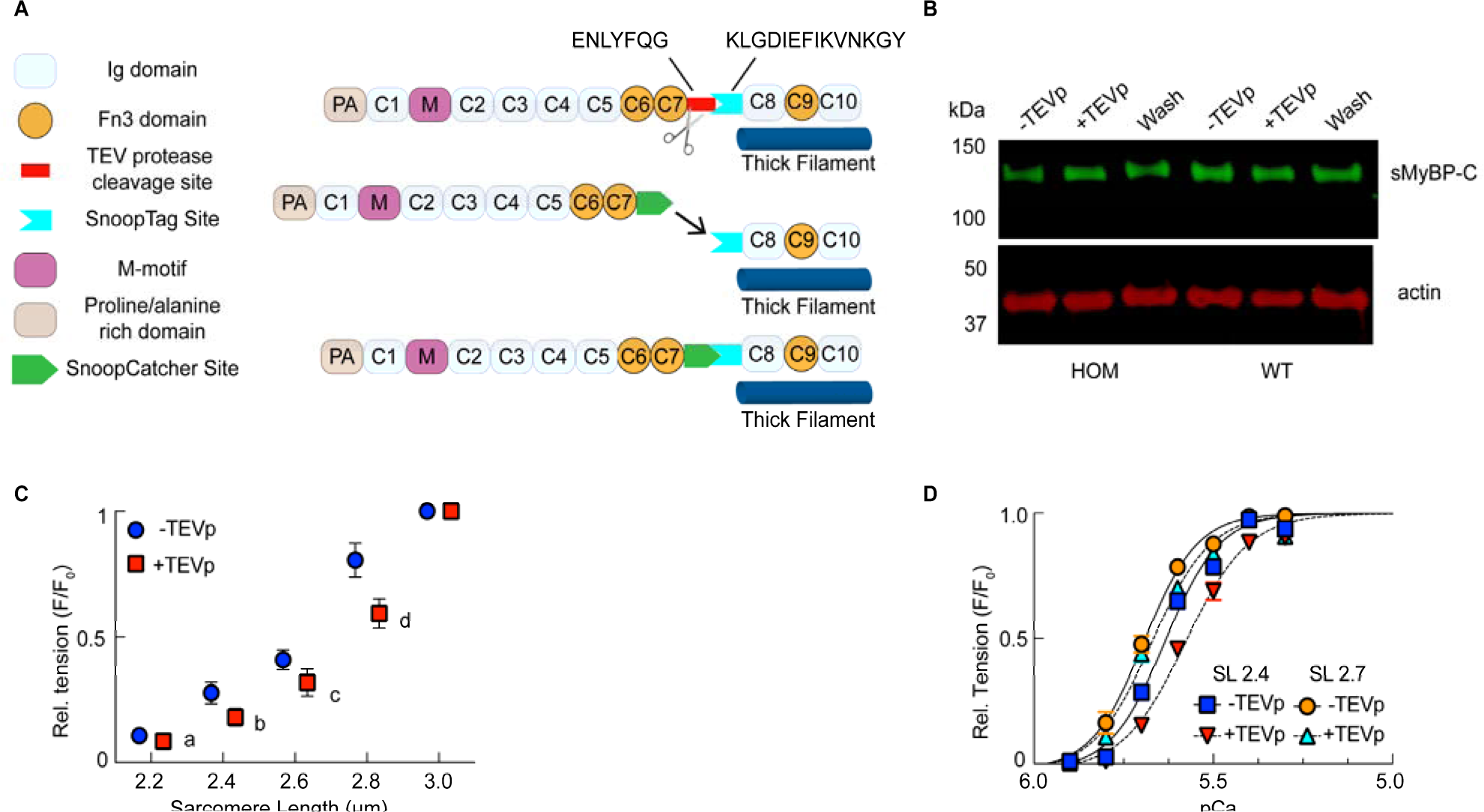
SNOOPC2 mouse line design and evaluation. (**A**) SNOOPC2 mice express a modified fMyBP-C that contains a TEV protease recognition site (red rectangle) and a SnoopTag (cyan trapezoid). Addition of TEV protease cleaves the endogenous C1-C7 domains of fMyBP-C while SnoopTag-C8-C10 remains anchored to the thick filament. Although not employed in this study, it is possible to incubate with a recombinant fMyBP-C construct containing the SnoopCatcher tag (green trapezoid), which will lead to *in situ* replacement of the cleaved fMyBP-C with the recombinant fragment. (**B**) Western blot of slow MyBP-C paralog from homozygous and wildtype SNOOPC2 psoas, before and after TEV protease treatment. As expected, no cleavage was detected in the slow MyBP-C paralog. (**C**) Relative passive tension-sarcomere length. (**D**) Relative tension-pCa relationship before and after cleavage, and at a short and long SL, with every condition scaled to its maximum (pCa 4.5) tension. Statistics throughout are repeated-measures ANOVA designs followed by a Tukey Honestly Significant Difference (HSD) post-hoc test on significant main effects. Different letters indicate differences after treatment at each SL. Data throughout reported as mean ± s.e.m. Further statistical details in Extended Data Table 1.

**Extended Data Figure 2.**
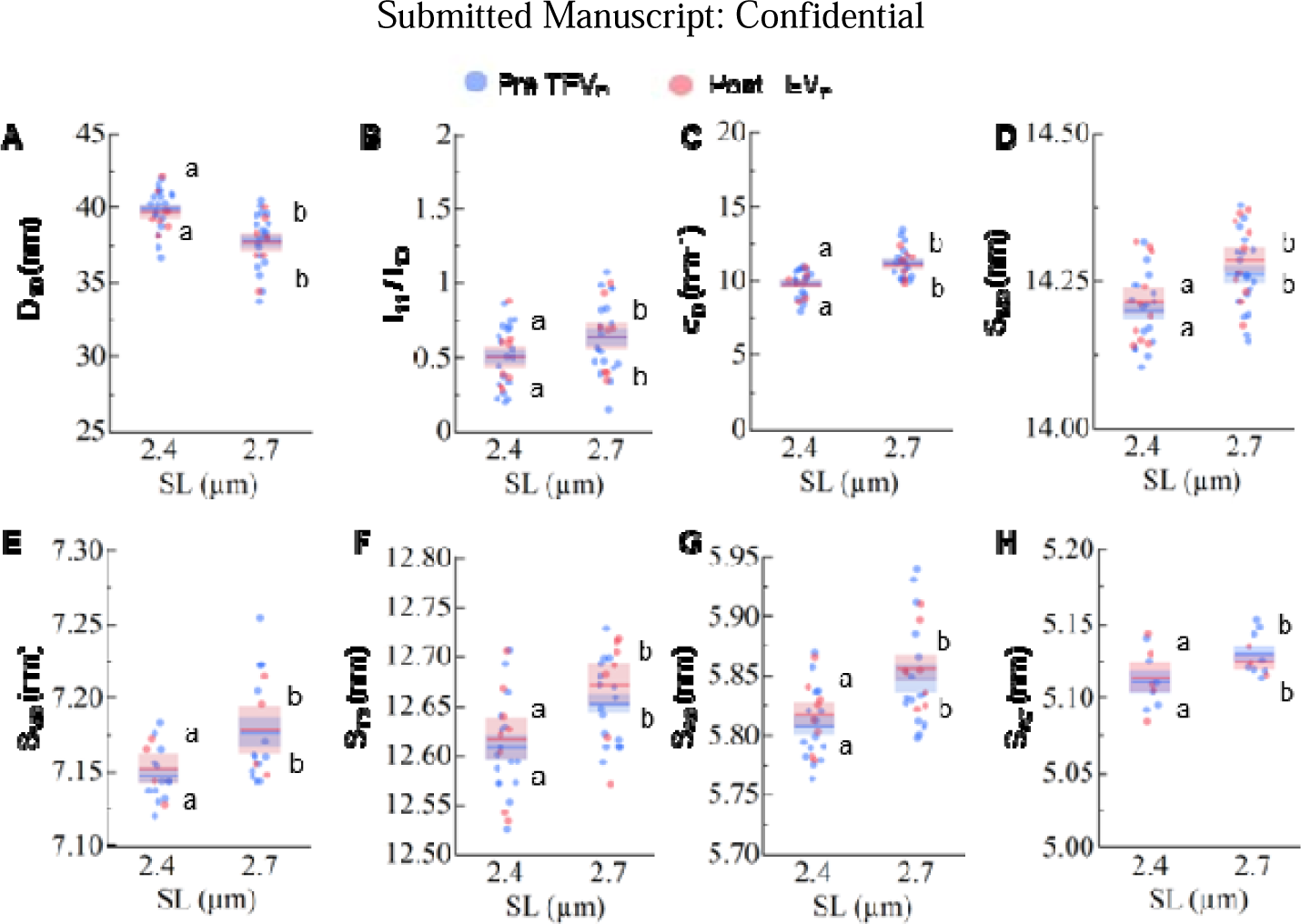
Control dataset for small-angle X-ray diffraction experiments. X-ray diffraction patterns were collected from wildtype SNOOPC2 psoas fibers under passive conditions at two difference sarcomere lengths (SL), before (blue) and after (red) TEV protease treatment. X-ray features shown for D_10_ (**A**), I_11_/I_10_ (**B**), σ_d_ (**C**), S_M3_ (**D**), S_M6_ (**E**), S_T3_ (**F**), S_A6_ (**G**), and S_A7_ (**H**). As expected, we detected no changes in structural features in wildtype muscles caused by TEV protease treatment. Statistics throughout are ANOVA designs with main effects treatment, SL, and their interaction, and a random effect of individual, followed by a Tukey Honestly Significant Difference (HSD) post-hoc test on significant main effects (P < 0.05), and reported in figures as connecting letters: different letters are significantly different. Data throughout reported as mean ± s.e.m. Full statistical details in Extended Data Table 7.

**Extended Data Table 1.** Statistical details from experiments shown in Fig. 1G-H and Extended Fig. 1D.

| Parameter | SL (μm) | Treatment | pCa | N | mean | s.e.m. | T-Test | df | Test Statistic | P-Value |
| --- | --- | --- | --- | --- | --- | --- | --- | --- | --- | --- |
| pCa50 | 2.4 | Pre | 4.5 - 9.0 | 9 | 5.61 | 0.02 | Treatment | 8 | T=8.54 | <0.0001 |
| pCa50 | 2.4 | Post | 4.5 - 9.0 | 9 | 5.55 | 0.02 |  |  |  |  |
| pCa50 | 2.7 | Pre | 4.5 - 9.0 | 9 | 5.70 | 0.01 | Treatment | 7 | T=1.86 | 0.11 |
| pCa50 | 2.7 | Post | 4.5 - 9.0 | 8 | 5.67 | 0.01 |  |  |  |  |
| Tn-pCa | 2.4 | Pre | 4.5 | 9 | 1.00 | 0.00 |  |  |  |  |
| Tn-pCa | 2.4 | Pre | 5.3 | 9 | 0.94 | 0.03 |  |  |  |  |
| Tn-pCa | 2.4 | Pre | 5.4 | 9 | 0.97 | 0.03 |  |  |  |  |
| Tn-pCa | 2.4 | Pre | 5.5 | 9 | 0.79 | 0.02 |  |  |  |  |
| Tn-pCa | 2.4 | Pre | 5.6 | 9 | 0.65 | 0.03 |  |  |  |  |
| Tn-pCa | 2.4 | Pre | 5.7 | 9 | 0.29 | 0.03 |  |  |  |  |
| Tn-pCa | 2.4 | Pre | 5.8 | 9 | 0.03 | 0.01 |  |  |  |  |
| Tn-pCa | 2.4 | Pre | 5.9 | 8 | 0.01 | 0.01 |  |  |  |  |
| Tn-pCa | 2.4 | Pre | 9 | 9 | 0.00 | 0.00 |  |  |  |  |
| Tn-pCa | 2.4 | Post | 4.5 | 9 | 1.00 | 0.00 |  |  |  |  |
| Tn-pCa | 2.4 | Post | 5.3 | 9 | 0.90 | 0.02 |  |  |  |  |
| Tn-pCa | 2.4 | Post | 5.4 | 9 | 0.88 | 0.02 |  |  |  |  |
| Tn-pCa | 2.4 | Post | 5.5 | 9 | 0.69 | 0.04 |  |  |  |  |
| Tn-pCa | 2.4 | Post | 5.6 | 9 | 0.46 | 0.02 |  |  |  |  |
| Tn-pCa | 2.4 | Post | 5.7 | 9 | 0.15 | 0.02 |  |  |  |  |
| Tn-pCa | 2.4 | Post | 5.8 | 9 | 0.01 | 0.01 |  |  |  |  |
| Tn-pCa | 2.4 | Post | 5.9 | 8 | 0.00 | 0.00 |  |  |  |  |
| Tn-pCa | 2.4 | Post | 9 | 8 | 0.00 | 0.00 |  |  |  |  |
| Tn-pCa | 2.7 | Pre | 4.5 | 9 | 1.00 | 0.00 |  |  |  |  |
| Tn-pCa | 2.7 | Pre | 5.4 | 9 | 0.99 | 0.02 |  |  |  |  |
| Tn-pCa | 2.7 | Pre | 5.5 | 9 | 0.88 | 0.02 |  |  |  |  |
| Tn-pCa | 2.7 | Pre | 5.6 | 9 | 0.79 | 0.03 |  |  |  |  |
| Tn-pCa | 2.7 | Pre | 5.7 | 9 | 0.48 | 0.03 |  |  |  |  |
| Tn-pCa | 2.7 | Pre | 5.8 | 9 | 0.16 | 0.04 |  |  |  |  |
| Tn-pCa | 2.7 | Pre | 5.9 | 9 | 0.01 | 0.01 |  |  |  |  |
| Tn-pCa | 2.7 | Pre | 9 | 9 | 0.00 | 0.00 |  |  |  |  |
| Tn-pCa | 2.7 | Post | 4.5 | 8 | 1.00 | 0.00 |  |  |  |  |
| Tn-pCa | 2.7 | Post | 5.4 | 8 | 0.97 | 0.02 |  |  |  |  |
| Tn-pCa | 2.7 | Post | 5.5 | 8 | 0.84 | 0.02 |  |  |  |  |
| Tn-pCa | 2.7 | Post | 5.6 | 8 | 0.70 | 0.02 |  |  |  |  |
| Tn-pCa | 2.7 | Post | 5.7 | 8 | 0.44 | 0.03 |  |  |  |  |
| Tn-pCa | 2.7 | Post | 5.8 | 8 | 0.11 | 0.03 |  |  |  |  |
| Tn-pCa | 2.7 | Post | 5.9 | 8 | 0.00 | 0.00 |  |  |  |  |
| Tn-pCa | 2.7 | Post | 9 | 8 | 0.00 | 0.00 |  |  |  |  |

Bold values indicate significant effect. s.e.m. = standard error of the mean

**Extended Data Table 2.** Statistical details from experiments shown in Fig. 1I, L, and M.

| Parameter | SL<br>($\mu\text{m}$) | Treatment | pCa | N | mean | s.e.m. | T-test | df | Test<br>Statistic | P-Value |
| --- | --- | --- | --- | --- | --- | --- | --- | --- | --- | --- |
| $K_{\text{tr}}$ | 2.4 | Pre | 4.5 | 9 | 1.00 | 0.00 | | | | |
| $K_{\text{tr}}$ | 2.4 | Post | 4.5 | 9 | 1.00 | 0.00 | | | | |
| $K_{\text{tr}}$ | 2.4 | Pre | 5.3 | 9 | 0.93 | 0.04 | Treatment | 8 | 1.80 | 0.110 |
| $K_{\text{tr}}$ | 2.4 | Post | 5.3 | 9 | 1.03 | 0.02 | | | | |
| $K_{\text{tr}}$ | 2.4 | Pre | 5.4 | 9 | 0.89 | 0.03 | Treatment | 8 | 1.94 | 0.088 |
| $K_{\text{tr}}$ | 2.4 | Post | 5.4 | 9 | 0.99 | 0.03 | | | | |
| $K_{\text{tr}}$ | 2.4 | Pre | 5.5 | 9 | 0.79 | 0.04 | Treatment | 8 | 4.19 | **0.003** |
| $K_{\text{tr}}$ | 2.4 | Post | 5.5 | 9 | 1.04 | 0.06 | | | | |
| $K_{\text{tr}}$ | 2.4 | Pre | 5.6 | 9 | 0.75 | 0.04 | Treatment | 8 | 4.21 | **0.003** |
| $K_{\text{tr}}$ | 2.4 | Post | 5.6 | 9 | 1.02 | 0.07 | | | | |
| $K_{\text{tr}}$ | 2.4 | Pre | 5.7 | 9 | 0.53 | 0.05 | Treatment | 8 | 1.04 | 0.330 |
| $K_{\text{tr}}$ | 2.4 | Post | 5.7 | 9 | 0.59 | 0.06 | | | | |
| $K_{\text{tr}}$ | 2.7 | Pre | 4.5 | 9 | 1.00 | 0.00 | | | | |
| $K_{\text{tr}}$ | 2.7 | Post | 4.5 | 8 | 1.00 | 0.00 | | | | |
| $K_{\text{tr}}$ | 2.7 | Pre | 5.4 | 9 | 0.93 | 0.02 | Treatment | 7 | 3.61 | **0.009** |
| $K_{\text{tr}}$ | 2.7 | Post | 5.4 | 8 | 1.05 | 0.02 | | | | |
| $K_{\text{tr}}$ | 2.7 | Pre | 5.5 | 9 | 0.91 | 0.02 | Treatment | 7 | 3.51 | **0.010** |
| $K_{\text{tr}}$ | 2.7 | Post | 5.5 | 8 | 1.06 | 0.04 | | | | |
| $K_{\text{tr}}$ | 2.7 | Pre | 5.6 | 9 | 0.91 | 0.04 | Treatment | 7 | 3.36 | **0.012** |
| $K_{\text{tr}}$ | 2.7 | Post | 5.6 | 8 | 1.07 | 0.05 | | | | |
| $K_{\text{tr}}$ | 2.7 | Pre | 5.7 | 8 | 0.69 | 0.05 | Treatment | 6 | 2.40 | 0.053 |
| $K_{\text{tr}}$ | 2.7 | Post | 5.7 | 8 | 0.79 | 0.06 | | | | |

Bold values indicate significant effect. s.e.m. = standard error of the mean. T-tests are one sided.

**Extended Data Table 3.** Statistical details from experiments shown in Fig. 1M.

| Parameter | SL ($\mu\text{m}$) | Treatment | pCa | N | mean | s.e.m. |
| --- | --- | --- | --- | --- | --- | --- |
| Abs. Tn-pCa | 2.4 | Pre | 4.5 | 9 | 26.74 | 4.95 |
| Abs. Tn-pCa | 2.4 | Pre | 5.3 | 9 | 23.31 | 3.96 |
| Abs. Tn-pCa | 2.4 | Pre | 5.4 | 9 | 24.02 | 4.28 |
| Abs. Tn-pCa | 2.4 | Pre | 5.5 | 9 | 20.10 | 3.45 |
| Abs. Tn-pCa | 2.4 | Pre | 5.6 | 9 | 15.90 | 2.80 |
| Abs. Tn-pCa | 2.4 | Pre | 5.7 | 9 | 7.84 | 2.02 |
| Abs. Tn-pCa | 2.4 | Pre | 5.8 | 9 | 1.21 | 0.63 |
| Abs. Tn-pCa | 2.4 | Pre | 5.9 | 8 | 0.36 | 0.25 |
| Abs. Tn-pCa | 2.4 | Pre | 9 | 9 | 0.00 | 0.00 |
| Abs. Tn-pCa | 2.4 | Post | 4.5 | 9 | 25.27 | 4.84 |
| Abs. Tn-pCa | 2.4 | Post | 5.3 | 9 | 22.56 | 4.23 |
| Abs. Tn-pCa | 2.4 | Post | 5.4 | 9 | 22.11 | 4.35 |
| Abs. Tn-pCa | 2.4 | Post | 5.5 | 9 | 18.40 | 4.03 |
| Abs. Tn-pCa | 2.4 | Post | 5.6 | 9 | 11.82 | 2.81 |
| Abs. Tn-pCa | 2.4 | Post | 5.7 | 9 | 4.17 | 1.23 |
| Abs. Tn-pCa | 2.4 | Post | 5.8 | 9 | 0.47 | 0.26 |
| Abs. Tn-pCa | 2.4 | Post | 5.9 | 8 | 0.06 | 0.05 |
| Abs. Tn-pCa | 2.4 | Post | 9 | 9 | 0.00 | 0.00 |
| Abs. Tn-pCa | 2.7 | Pre | 4.5 | 9 | 49.48 | 7.31 |
| Abs. Tn-pCa | 2.7 | Pre | 5.4 | 9 | 45.13 | 7.00 |
| Abs. Tn-pCa | 2.7 | Pre | 5.5 | 9 | 42.35 | 7.17 |
| Abs. Tn-pCa | 2.7 | Pre | 5.6 | 9 | 35.97 | 6.22 |
| Abs. Tn-pCa | 2.7 | Pre | 5.7 | 9 | 24.12 | 4.89 |
| Abs. Tn-pCa | 2.7 | Pre | 5.8 | 9 | 9.55 | 4.01 |
| Abs. Tn-pCa | 2.7 | Pre | 5.9 | 9 | 0.51 | 0.43 |
| Abs. Tn-pCa | 2.7 | Pre | 9 | 9 | 0.00 | 0.00 |
| Abs. Tn-pCa | 2.7 | Post | 4.5 | 8 | 57.22 | 8.90 |
| Abs. Tn-pCa | 2.7 | Post | 5.4 | 8 | 51.57 | 6.38 |
| Abs. Tn-pCa | 2.7 | Post | 5.5 | 8 | 47.20 | 7.59 |
| Abs. Tn-pCa | 2.7 | Post | 5.6 | 8 | 37.44 | 4.88 |
| Abs. Tn-pCa | 2.7 | Post | 5.7 | 8 | 25.15 | 4.04 |
| Abs. Tn-pCa | 2.7 | Post | 5.8 | 8 | 6.18 | 1.45 |
| Abs. Tn-pCa | 2.7 | Post | 5.9 | 8 | 0.25 | 0.20 |
| Abs. Tn-pCa | 2.7 | Post | 9 | 8 | 0.00 | 0.00 |

s.e.m. = standard error of the mean

**Extended Data Table 4.** Statistical details from experiments shown in Figure 1F and Extended Figure 1C.

| Parameter | SL ($\mu\text{m}$) | Treatment | N | mean | s.e.m. | ANOVA | DF<sub>1</sub> | DF<sub>2</sub> | F stat | Prob > F |
| --- | --- | --- | --- | --- | --- | --- | --- | --- | --- | --- |
| Abs. Passive Force | 2.2 | Pre | 9 | 0.92 | 0.06 | SL | 4 | 85 | 0.75 | <0.0001 |
| Abs. Passive Force | 2.2 | Post | 10 | 0.87 |  | Treatment | 4 | 85 | 35.82 | 0.99 |
| Abs. Passive Force | 2.4 | Pre | 9 | 2.38 | 0.28 | SL*Treatment | 1 | 85 | 0.00024 | 0.56 |
| Abs. Passive Force | 2.4 | Post | 10 | 1.82 | 0.18 |  |  |  |  |  |
| Abs. Passive Force | 2.6 | Pre | 9 | 3.75 | 0.34 |  |  |  |  |  |
| Abs. Passive Force | 2.6 | Post | 10 | 3.39 | 0.34 |  |  |  |  |  |
| Abs. Passive Force | 2.8 | Pre | 9 | 7.72 | 1.04 |  |  |  |  |  |
| Abs. Passive Force | 2.8 | Post | 10 | 6.64 | 0.80 |  |  |  |  |  |
| Abs. Passive Force | 3 | Pre | 9 | 10.27 | 1.70 |  |  |  |  |  |
| Abs. Passive Force | 3 | Post | 10 | 12.37 | 2.21 |  |  |  |  |  |
| Norm. Passive Force | 2.2 | Pre | 9 | 0.11 | 0.02 | SL | 3 | 68 | 1.79 | <0.0001 |
| Norm. Passive Force | 2.2 | Post | 10 | 0.09 | 0.01 | Treatment | 3 | 68 | 81.76 | 0.0007 |
| Norm. Passive Force | 2.4 | Pre | 9 | 0.28 | 0.04 | SL*Treatment | 1 | 68 | 12.51 | 0.16 |
| Norm. Passive Force | 2.4 | Post | 10 | 0.18 | 0.03 |  |  |  |  |  |
| Norm. Passive Force | 2.6 | Pre | 9 | 0.41 | 0.04 |  |  |  |  |  |
| Norm. Passive Force | 2.6 | Post | 10 | 0.32 | 0.04 |  |  |  |  |  |
| Norm. Passive Force | 2.8 | Pre | 9 | 0.81 | 0.07 |  |  |  |  |  |
| Norm. Passive Force | 2.8 | Post | 10 | 0.60 | 0.05 |  |  |  |  |  |
| Norm. Passive Force | 3 | Pre | 9 | 1.00 | 0.00 |  |  |  |  |  |
| Norm. Passive Force | 3 | Post | 10 | 1.00 | 0.00 |  |  |  |  |  |

Bold values indicate significant effect. s.e.m. = standard error of the mean

**Extended Data Table 5.** Statistical details from experiments shown in Figure 2.

| Parameter | SL ($\mu\text{m}$) | Treatment | N | mean | s.e.m. | ANOVA | DF<sub>1</sub> | DF<sub>2</sub> | F stat | Prob > F |
| --- | --- | --- | --- | --- | --- | --- | --- | --- | --- | --- |
| d<sub>10</sub> (nm) | 2.40 | Pre | 36 | 39.004 | 0.363 | SL | 2 | 162 | 88.07 | < **0.0001** |
| d<sub>10</sub> (nm) | 2.40 | Post | 33 | 39.763 | 0.313 | Treatment | 1 | 163 | 34.55 | < **0.0001** |
| d<sub>10</sub> (nm) | 2.70 | Pre | 33 | 37.912 | 0.441 | Interaction | 2 | 162 | 0.08 | 0.92 |
| d<sub>10</sub> (nm) | 2.70 | Post | 34 | 38.761 | 0.343 |  |  |  |  |  |
| d<sub>10</sub> (nm) | 3.00 | Pre | 32 | 36.646 | 0.487 |  |  |  |  |  |
| d<sub>10</sub> (nm) | 3.00 | Post | 34 | 37.361 | 0.426 |  |  |  |  |  |
| $\sigma_D$ (nm<sup>-1</sup>) | 2.40 | Pre | 21 | 7.817 | 0.253 | SL | 2 | 86 | 39.58 | < **0.0001** |
| $\sigma_D$ (nm<sup>-1</sup>) | 2.40 | Post | 20 | 8.103 | 0.304 | Treatment | 1 | 88 | 0.11 | 0.74 |
| $\sigma_D$ (nm<sup>-1</sup>) | 2.70 | Pre | 18 | 9.226 | 0.426 | Interaction | 2 | 86 | 0.50 | 0.61 |
| $\sigma_D$ (nm<sup>-1</sup>) | 2.70 | Post | 18 | 8.902 | 0.424 | | | | | |
| $\sigma_D$ (nm<sup>-1</sup>) | 3.00 | Pre | 15 | 10.293 | 0.603 | | | | | |
| $\sigma_D$ (nm<sup>-1</sup>) | 3.00 | Post | 21 | 10.616 | 0.524 | | | | | |
| S<sub>T3</sub> (nm) | 2.40 | Pre | 33 | 12.644 | 0.007 | SL | 2 | 144 | 7.48 | < **0.001** |
| S<sub>T3</sub> (nm) | 2.40 | Post | 29 | 12.631 | 0.007 | Treatment | 1 | 147 | 13.73 | < **0.001** |
| S<sub>T3</sub> (nm) | 2.70 | Pre | 29 | 12.650 | 0.007 | Interaction | 2 | 144 | 0.03 | 0.97 |
| S<sub>T3</sub> (nm) | 2.70 | Post | 31 | 12.638 | 0.009 |  |  |  |  |  |
| S<sub>T3</sub> (nm) | 3.00 | Pre | 29 | 12.659 | 0.007 |  |  |  |  |  |
| S<sub>T3</sub> (nm) | 3.00 | Post | 32 | 12.647 | 0.007 |  |  |  |  |  |
| S<sub>A6</sub> (nm) | 2.40 | Pre | 32 | 5.818 | 0.005 | SL | 2 | 134 | 54.02 | < **0.0001** |
| S<sub>A6</sub> (nm) | 2.40 | Post | 24 | 5.804 | 0.005 | Treatment | 1 | 136 | 36.81 | < **0.0001** |
| S<sub>A6</sub> (nm) | 2.70 | Pre | 30 | 5.841 | 0.005 | SL*Treatment | 2 | 134 | 1.12 | 0.33 |
| S<sub>A6</sub> (nm) | 2.70 | Post | 27 | 5.826 | 0.005 |  |  |  |  |  |
| S<sub>A6</sub> (nm) | 3.00 | Pre | 29 | 5.845 | 0.005 |  |  |  |  |  |
| S<sub>A6</sub> (nm) | 3.00 | Post | 29 | 5.835 | 0.005 |  |  |  |  |  |
| S<sub>A7</sub> (nm) | 2.40 | Pre | 28 | 5.062 | 0.003 | SL | 2 | 110 | 26.01 | < **0.0001** |
| S<sub>A7</sub> (nm) | 2.40 | Post | 23 | 5.046 | 0.004 | Treatment | 1 | 113 | 15.42 | < **0.0001** |
| S<sub>A7</sub> (nm) | 2.70 | Pre | 22 | 5.071 | 0.004 | SL*Treatment | 2 | 109 | 2.35 | 0.10 |
| S<sub>A7</sub> (nm) | 2.70 | Post | 20 | 5.060 | 0.004 |  |  |  |  |  |
| S<sub>A7</sub> (nm) | 3.00 | Pre | 26 | 5.076 | 0.003 |  |  |  |  |  |
| S<sub>A7</sub> (nm) | 3.00 | Post | 23 | 5.073 | 0.004 |  |  |  |  |  |
| S<sub>gActin</sub> (nm) | 2.40 | Pre | 28 | 2.708 | 0.002 | SL | 2 | 99 | 65.67 | < **0.0001** |
| S<sub>gActin</sub> (nm) | 2.40 | Post | 20 | 2.699 | 0.002 | Treatment | 1 | 101 | 46.77 | < **0.0001** |
| S<sub>gActin</sub> (nm) | 2.70 | Pre | 22 | 2.715 | 0.002 | SL*Treatment | 2 | 99 | 3.62 | **0.03** |
| S<sub>gActin</sub> (nm) | 2.70 | Post | 17 | 2.709 | 0.002 |  |  |  |  |  |
| S<sub>gActin</sub> (nm) | 3.00 | Pre | 24 | 2.718 | 0.002 |  |  |  |  |  |
| S<sub>gActin</sub> (nm) | 3.00 | Post | 22 | 2.715 | 0.002 |  |  |  |  |  |

Bold values indicate significant effect. s.e.m. = standard error of the mean

**Extended Data Table 6.** Statistical details from experiments shown in Figure 3.

| Parameter | SL ($\mu\text{m}$) | Treatment | N | mean | s.e.m. | ANOVA | DF<sub>1</sub> | DF<sub>2</sub> | F stat | Prob > F |
| --- | --- | --- | --- | --- | --- | --- | --- | --- | --- | --- |
| I<sub>M2</sub> cluster | 2.40 | Pre | 28 | 0.371 | 0.036 | SL | 2 | 129 | 4.05 | **0.02** |
| I<sub>M2</sub> cluster | 2.40 | Post | 28 | 0.206 | 0.022 | Treatment | 1 | 135 | 26.60 | < **0.0001** |
| I<sub>M2</sub> cluster | 2.70 | Pre | 24 | 0.292 | 0.035 | Interaction | 2 | 129 | 4.91 | **0.01** |
| I<sub>M2</sub> cluster | 2.70 | Post | 27 | 0.216 | 0.019 |  |  |  |  |  |
| I<sub>M2</sub> cluster | 3.00 | Pre | 28 | 0.242 | 0.021 |  |  |  |  |  |
| I<sub>M2</sub> cluster | 3.00 | Post | 28 | 0.211 | 0.022 |  |  |  |  |  |
| I<sub>11</sub>/I<sub>10</sub> | 2.40 | Pre | 35 | 0.636 | 0.029 | SL | 2 | 157 | 4.67 | **0.01** |
| I<sub>11</sub>/I<sub>10</sub> | 2.40 | Post | 31 | 0.749 | 0.028 | Treatment | 1 | 159 | 19.42 | < **0.0001** |
| I<sub>11</sub>/I<sub>10</sub> | 2.70 | Pre | 32 | 0.726 | 0.042 | Interaction | 2 | 157 | 0.35 | 0.71 |
| I<sub>11</sub>/I<sub>10</sub> | 2.70 | Post | 34 | 0.827 | 0.035 |  |  |  |  |  |
| I<sub>11</sub>/I<sub>10</sub> | 3.00 | Pre | 30 | 0.708 | 0.043 |  |  |  |  |  |
| I<sub>11</sub>/I<sub>10</sub> | 3.00 | Post | 33 | 0.860 | 0.053 |  |  |  |  |  |
| S<sub>M3</sub> (nm) | 2.40 | Pre | 32 | 14.184 | 0.008 | SL | 2 | 146 | 36.34 | < **0.0001** |
| S<sub>M3</sub> (nm) | 2.40 | Post | 28 | 14.223 | 0.007 | Treatment | 1 | 147 | 18.24 | < **0.0001** |
| S<sub>M3</sub> (nm) | 2.70 | Pre | 31 | 14.231 | 0.009 | Interaction | 2 | 143 | 3.39 | **0.04** |
| S<sub>M3</sub> (nm) | 2.70 | Post | 31 | 14.264 | 0.009 |  |  |  |  |  |
| S<sub>M3</sub> (nm) | 3.00 | Pre | 31 | 14.259 | 0.009 |  |  |  |  |  |
| S<sub>M3</sub> (nm) | 3.00 | Post | 31 | 14.266 | 0.010 |  |  |  |  |  |
| S<sub>M6</sub> (nm) | 2.40 | Pre | 31 | 7.163 | 0.002 | SL | 2 | 134 | 30.07 | < **0.0001** |
| S<sub>M6</sub> (nm) | 2.40 | Post | 26 | 7.169 | 0.002 | Treatment | 1 | 137 | 12.75 | < **0.001** |
| S<sub>M6</sub> (nm) | 2.70 | Pre | 29 | 7.171 | 0.002 | Interaction | 2 | 134 | 0.12 | 0.89 |
| S<sub>M6</sub> (nm) | 2.70 | Post | 29 | 7.176 | 0.003 |  |  |  |  |  |
| S<sub>M6</sub> (nm) | 3.00 | Pre | 28 | 7.176 | 0.003 |  |  |  |  |  |
| S<sub>M6</sub> (nm) | 3.00 | Post | 27 | 7.180 | 0.003 |  |  |  |  |  |

Bold values indicate significant effect. s.e.m. = standard error of the mean

**Extended Data Table 7.** Statistical details from experiments shown Extended Data Figure 2 (controls)

| Parameter | SL ($\mu\text{m}$) | Treatment | N | mean | s.e.m. | ANOVA | DF<sub>1</sub> | DF<sub>2</sub> | F stat | Prob > F |
| --- | --- | --- | --- | --- | --- | --- | --- | --- | --- | --- |
| d<sub>10</sub> (nm) | 2.4 | Pre | 19 | 39.938 | 0.325 | Treatment | 1 | 37 | 0.73 | 0.40 |
| d<sub>10</sub> (nm) | 2.4 | Post | 8 | 39.740 | 0.460 | SL | 1 | 33 | 48.00 | **< 0.001** |
| d<sub>10</sub> (nm) | 2.7 | Pre | 19 | 37.774 | 0.441 | Interaction | 1 | 33 | 0.04 | 0.83 |
| d<sub>10</sub> (nm) | 2.7 | Post | 8 | 37.702 | 0.621 |  |  |  |  |  |
| I<sub>11</sub>/I<sub>10</sub> | 2.4 | Pre | 19 | 0.504 | 0.049 | Treatment | 1 | 37 | 0.10 | 0.75 |
| I<sub>11</sub>/I<sub>10</sub> | 2.4 | Post | 8 | 0.501 | 0.073 | SL | 1 | 32 | 9.20 | **<0.01** |
| I<sub>11</sub>/I<sub>10</sub> | 2.7 | Pre | 19 | 0.634 | 0.058 | Interaction | 1 | 32 | 0.01 | 0.91 |
| I<sub>11</sub>/I<sub>10</sub> | 2.7 | Post | 8 | 0.642 | 0.088 |  |  |  |  |  |
| $\sigma_D$ (nm<sup>-1</sup>) | 2.4 | Pre | 17 | 9.838 | 0.237 | Treatment | 1 | 45 | 0.23 | 0.63 |
| $\sigma_D$ (nm<sup>-1</sup>) | 2.4 | Post | 7 | 9.771 | 0.316 | SL | 1 | 28 | 21.38 | **< 0.001** |
| $\sigma_D$ (nm<sup>-1</sup>) | 2.7 | Pre | 17 | 11.220 | 0.262 | Interaction | 1 | 26 | 0.00 | 0.96 |
| $\sigma_D$ (nm<sup>-1</sup>) | 2.7 | Post | 8 | 11.107 | 0.306 | | | | | |
| S<sub>M3</sub> (nm) | 2.4 | Pre | 17 | 14.200 | 0.015 | Treatment | 1 | 37 | 0.02 | 0.88 |
| S<sub>M3</sub> (nm) | 2.4 | Post | 10 | 14.216 | 0.022 | SL | 1 | 33 | 29.96 | **< 0.001** |
| S<sub>M3</sub> (nm) | 2.7 | Pre | 19 | 14.261 | 0.015 | Treatment*SL | 1 | 33 | 0.08 | 0.78 |
| S<sub>M3</sub> (nm) | 2.7 | Post | 10 | 14.286 | 0.022 |  |  |  |  |  |
| S<sub>M6</sub> (nm) | 2.4 | Pre | 13 | 7.148 | 0.005 | Treatment | 1 | 23 | 0.04 | 0.83 |
| S<sub>M6</sub> (nm) | 2.4 | Post | 4 | 7.152 | 0.010 | SL | 1 | 18 | 13.38 | **<0.01** |
| S<sub>M6</sub> (nm) | 2.7 | Pre | 13 | 7.177 | 0.010 | Treatment*SL | 1 | 17 | 0.02 | 0.88 |
| S<sub>M6</sub> (nm) | 2.7 | Post | 4 | 7.179 | 0.016 |  |  |  |  |  |
| S<sub>T3</sub> (nm) | 2.4 | Pre | 16 | 12.608 | 0.012 | Treatment | 1 | 32 | 0.29 | 0.59 |
| S<sub>T3</sub> (nm) | 2.4 | Post | 8 | 12.617 | 0.021 | SL | 1 | 26 | 27.00 | **< 0.001** |
| S<sub>T3</sub> (nm) | 2.7 | Pre | 16 | 12.652 | 0.010 | Treatment*SL | 1 | 26 | 0.01 | 0.91 |
| S<sub>T3</sub> (nm) | 2.7 | Post | 7 | 12.671 | 0.021 |  |  |  |  |  |
| S<sub>A6</sub> (nm) | 2.4 | Pre | 15 | 5.808 | 0.008 | Treatment | 1 | 28 | 1.41 | 0.24 |
| S<sub>A6</sub> (nm) | 2.4 | Post | 8 | 5.817 | 0.010 | SL | 1 | 26 | 51.34 | **< 0.001** |
| S<sub>A6</sub> (nm) | 2.7 | Pre | 16 | 5.847 | 0.011 | Treatment*SL | 1 | 26 | 0.22 | 0.64 |
| S<sub>A6</sub> (nm) | 2.7 | Post | 8 | 5.856 | 0.011 |  |  |  |  |  |
| S<sub>A7</sub> (nm) | 2.4 | Pre | 6 | 5.111 | 0.008 | Treatment | 1 | 17 | 0.15 | 0.70 |
| S<sub>A7</sub> (nm) | 2.4 | Post | 5 | 5.114 | 0.010 | SL | 1 | 11 | 6.26 | **0.03** |
| S<sub>A7</sub> (nm) | 2.7 | Pre | 8 | 5.130 | 0.005 | Treatment*SL | 1 | 10 | 0.11 | 0.74 |
| S<sub>A7</sub> (nm) | 2.7 | Post | 3 | 5.125 | 0.006 |  |  |  |  |  |

Bold values indicate significant effect. s.e.m. = standard error of the mean

